# Is developmental plasticity triggered by DNA methylation changes in the invasive cane toad (*Rhinella marina*)?

**DOI:** 10.1101/2023.10.26.564158

**Authors:** Boris Yagound, Roshmi R. Sarma, Richard J. Edwards, Mark F. Richardson, Carlos M. Rodriguez Lopez, Michael R. Crossland, Gregory P. Brown, Jayna L. DeVore, Richard Shine, Lee A. Rollins

**Author notes:** Current institution (Roshmi R. Sarma, Richard J. Edwards, Carlos M. Rodriguez Lopez, Gregory P. Brown, Jayna L. DeVore, Richard Shine, Lee A. Rollins). Corresponding author: Boris Yagound.

## Abstract

Many organisms can adjust their development according to environmental conditions, including the presence of conspecifics. Although this developmental plasticity is common in amphibians, its underlying molecular mechanisms remain largely unknown. Exposure during development to either ‘cannibal cues’ from older conspecifics, or ‘alarm cues’ from injured conspecifics, causes reduced growth and survival in cane toad (*Rhinella marina*) tadpoles. Epigenetic modifications, such as changes in DNA methylation patterns, are a plausible mechanism underlying these developmental plastic responses. Here we tested this hypothesis, and asked whether cannibal cues and alarm cues trigger the same DNA methylation changes in developing cane toads. We found that exposure to both cannibal cues and alarm cues induced local changes in DNA methylation patterns. These DNA methylation changes affected genes putatively involved in developmental processes, but in different genomic regions for different conspecific-derived cues. Genetic background explained most of the epigenetic variation among individuals. Overall, the molecular mechanisms triggered by exposure to cannibal cues seem to differ from those triggered by alarm cues. Studies linking epigenetic modifications to transcriptional activity are needed to clarify the proximate mechanisms that regulate developmental plasticity in cane toads.

## Introduction

Developmental plasticity is the ability for a single genotype to give rise to a range of phenotypes in different environments. Plasticity can be adaptive when environmental conditions are predictable, and can involve both short- and long-term changes in physiology, morphology, life-history traits and behaviour (Sultan, 2003; West-Eberhard, 2003). Although studies on the underlying molecular mechanisms of plasticity are burgeoning, these mechanisms remain poorly understood (Gilbert & Epel, 2015; Lafuente & Beldade, 2019; Sommer, 2020).

Amphibians are good models to study molecular mechanisms of phenotypic plasticity because the growth and development of their aquatic larvae are susceptible to environmental factors such as food availability, predator exposure, conspecific density, and pond drying (Denver, 2021; Newman, 1992; Wilbur & Collins, 1973). The invasive cane toad, *Rhinella marina*, is a good exemplar of a species capable of altering its developmental trajectory in response to environmental conditions (Lever, 2001). Female cane toads lay large clutches (up to several tens of thousands of eggs) asynchronously in ponds, resulting in overlapping cohorts of developing offspring where egg and larvae densities can reach extreme levels (Alford, Cohen, Crossland, Hearnden, & Schwarzkopf, 1995; DeVore, Crossland, & Shine, 2021). Cannibalism of eggs and hatchlings by older tadpoles is common in these breeding ponds, such that survival of newly-laid eggs to the free-swimming tadpole stage is often <1% (Alford et al., 1995; DeVore, Crossland, & Shine, 2021).

Interestingly, the presence of conspecifics affects cane toad larval development in two contexts (Crossland & Shine, 2012; Hagman, Hayes, Capon, & Shine, 2009). First, hatchlings are affected by ‘cannibal cues’ associated with the approach of older, cannibalistic tadpoles (Clarke, Crossland, Shilton, Shine, & Rohr, 2015; Crossland & Shine, 2011, 2012; DeVore, Crossland, & Shine, 2021). Exposure to cannibal cues causes hatchlings to accelerate development, with significant carry-over effects during the subsequent tadpole stage: decreased tadpole survival, decreased body mass and body size (i.e., growth), reduced tooth row keratinization, increased swimming behaviour and repression of feeding behaviour (Clarke et al., 2015; Crossland & Shine, 2012; DeVore, Crossland, & Shine, 2021; DeVore, Crossland, Shine, & Ducatez, 2021; McCann, Crossland, Greenlees, & Shine, 2020). Second, injured tadpoles release an ‘alarm cue’, reflecting a predation risk, that elicits immediate avoidance by conspecifics (Hagman & Shine, 2008). Chronic exposure to alarm cues during tadpole development decreases tadpole survival, reduces growth at metamorphosis, increases the size of parotoid glands, and can reduce development rate (Crossland, Salim, Capon, & Shine, 2019; Hagman et al., 2009; Hagman & Shine, 2009).

Both alarm cues and cannibal cues thus induce developmental plastic responses and reduce growth and survival in cane toad tadpoles. Moreover, both cues likely have long-term detrimental effects post metamorphosis, affecting adult survival against cannibalism (Pizzatto & Shine, 2008), predation (Ward-Fear, Brown, & Shine, 2012), parasitism (Kelehear, Webb, & Shine, 2009) and desiccation (Child, Phillips, & Shine, 2008). Finally, these cues appear mechanistically associated with one another, because exposure to alarm cues can have intergenerational effects by increasing the potency of cannibal cues in the next generation (Sarma et al., 2021).

In this study, we asked whether exposure to cannibal cues and exposure to alarm cues might trigger the same molecular mechanisms. Specifically, we tested i) whether exposure to conspecific cues triggers changes in DNA methylation patterns in cane toad tadpoles that might then induce developmental plasticity, and ii) whether both cannibal cues and alarm cues trigger the same epigenetic modifications. We focused on DNA methylation, because this epigenetic mechanism can influence transcriptional activity on the one hand (Jaenisch & Bird, 2003), and can be affected by environmental factors on the other (Dowen et al., 2012;

Radford et al., 2014), and because studies of cane toads have shown that exposure to alarm cues changes DNA methylation patterns in tadpoles (Sarma et al., 2021; Sarma et al., 2020). Changes in DNA methylation are thus a plausible molecular mechanism that might underlie developmental plasticity in cane toads.

## Material and Methods

### Toads

We collected adult cane toads from two genetically-distinct populations (Selechnik et al., 2019) within the Australian invasive range: ‘range core’ and ‘range edge’. We collected range-core toads from five localities in Queensland: Townsville, Mission Beach, Port Douglas, Innisfail and Tully. We collected range-edge toads from one locality in the Northern Territory, Middle Point, and from four localities in Western Australia: Doongan, Lake Argyle, Mary Pool and Oombulgarri (Figure S1). We transported all collected toads to our field station in Middle Point, Northern Territory, where they were maintained in outdoor enclosures with refugia, water and a constant food supply. We subcutaneously injected two pairs of toads from each locality with synthetic gonadotropin to induce spawning (see DeVore, Crossland, and Shine (2021) for details). We then kept eggs in aerated holding tanks for 72 h to ensure successful fertilisation.

### Experimental design

In this study, we compared the effect of conspecific exposure on DNA methylation during development in two contexts: exposure to cannibal cues and exposure to alarm cues.

### Cannibal cue experiment

In this experiment, we exposed focal hatchlings from eight clutches (i.e., 4 range-core and 4 range-edge clutches) to three treatments: 1) exposure to conspecific tadpoles (i.e., cannibal cues) from clutch *i*, 2) exposure to conspecific tadpoles from clutch *j*, 3) control (no conspecific tadpoles added). Clutches *i* and *j* were raised from clutches from the same locality as the focal hatchlings but were included so that we could measure the impact of genotype on the effect of cannibal cues. We used 1l-treatment tanks filled with 750 ml of water from a local aquifer. In the cannibal cue treatments, we first added three captive-raised tadpoles [developmental stage 30–35, Gosner (1960)] to each tank, separated by a 1×1 mm fly screen mesh from the compartment where focal hatchlings were placed. This allowed chemical cannibal cues to diffuse throughout the container but prevented cannibalism (Crossland & Shine, 2012). Several hours after conspecific tadpoles were added, we randomly allocated five hatchlings [developmental stage 16/17, i.e., approximately one day post-hatching, Gosner (1960)] to each treatment tank. We replicated each treatment three times. We removed the conspecific tadpoles after 24 h and left the developing hatchlings for a further 24 h, by which time they had developed into free-swimming, feeding tadpoles (stage 25). We then transferred the stage 25 tadpoles to new tanks, and fed them blended Hikari algae wafers (Kyorin, Japan) ad libitum, with fresh water changes every three days. Ten days later, we humanely euthanised all tadpoles (stage ∼32) and three tadpoles from each tank were blotted, weighed, and then frozen prior to DNA extraction.

### Alarm cue experiment

This experiment was described in Sarma et al. (2020). Briefly, we randomly allocated two hatchlings from each clutch to two treatments: 1) exposure to conspecific alarm cues (*n* = 11 clutches, each region), 2) control (*n* = 10 clutches, range core; *n* = 9 clutches, range edge). We used hatchlings from the same clutches for each of these two treatments, unless mortality prevented us from doing so (i.e., three cases where only one hatchling was available and was allocated to the alarm cue treatment). In the alarm cue treatment, we added to each tank 4 ml of water containing the freshly-macerated bodies of two conspecific tadpoles (Hagman et al., 2009) on each of ten consecutive days (days 7–16 post spawning). In the control treatment, we added 4 ml of non-chlorinated water on ten consecutive days. Two days later, we humanely euthanised two 18-days old tadpoles (stage ∼35) per tank, weighed them, and preserved them in RNALater; one of these individuals was used for DNA methylation analysis.

### Reduced-representation bisulfite sequencing

We extracted DNA from whole tadpoles using a Gentra Puregene Kit (Qiagen) according to the manufacturer’s instructions. We prepared reduced-representation bisulfite sequencing (RRBS) libraries using 100 ng of genomic DNA per sample with the Ovation RRBS Methyl-Seq Kit (NuGEN Technologies, San Carlos, USA). Libraries (100 bp single-end) were sequenced on a NovaSeq 6000 S2 flowcell platform (Illumina, San Diego, USA). RRBS library preparation and sequencing were conducted at the Ramaciotti Centre for Genomics (UNSW, Sydney, Australia).

### DNA methylation analyses

We used FastQC 0.11.8 (Andrews, 2010) to assess the quality of the reads. We trimmed adapter sequences and low-quality reads using Trim Galore 6.5 (Krueger, 2015). We then mapped the remaining reads to the cane toad genome (Edwards et al., 2018) using Bismark 0.22.3 (Krueger & Andrews, 2011) with HISAT2 2.1.0 (Kim, Paggi, Park, Bennett, & Salzberg, 2019). We extracted methylation status for each CpG using Bismark. We carried out downstream differential methylation analyses using the package methylKit (Akalin et al., 2012) in R 4.0.4 (R Core Team, 2021). We merged strands for each CpG, and we only kept sites that had a depth of coverage of at least 10 reads for subsequent analyses. We further filtered out any site with a coverage higher than the 99.9th percentile of read counts. We defined differentially methylated cytosines (hereafter, DMCs) as CpGs with a methylation difference of 20% or greater between the two groups (i.e., conspecific-cue exposed vs controls), a *q*-value (Fisher’s exact test corrected *p*-value) (Wang, Tuominen, & Tsai, 2011) of 0.05 or less, and that were present in at least three samples in each group. We used the R package eDMR (Li et al., 2013) to identify differentially methylated regions (hereafter, DMRs). DMRs were defined as regions with a mean methylation difference of at least 20% between the two groups, a *q*-value of 0.05 or less, and that contained at least 5 CpGs and 3 DMCs.

### Effect of conspecific cues on tadpole growth

For each experiment, we investigated whether the exposure to conspecific cues influenced mass of focal tadpoles, using linear mixed effects models (LMMs) with the R package lme4 (Bates, Machler, Bolker, & Walker, 2015). We included mass as the dependent variable, treatment as a fixed effect, and clutch ID and replicate as random factors.

## Results

The average mass of tadpoles that had been exposed to cannibal cues as hatchlings was lower than controls in range-edge populations, but not in range-core populations (LMMs, respectively *p* < 0.00001 and *p* = 0.129; Figure 1 A–B). Alarm cues had no significant effect on mean mass of exposed tadpoles in either range-core or range-edge populations (both *p* > 0.396; Figure 1 C–D).

**Figure 1.**
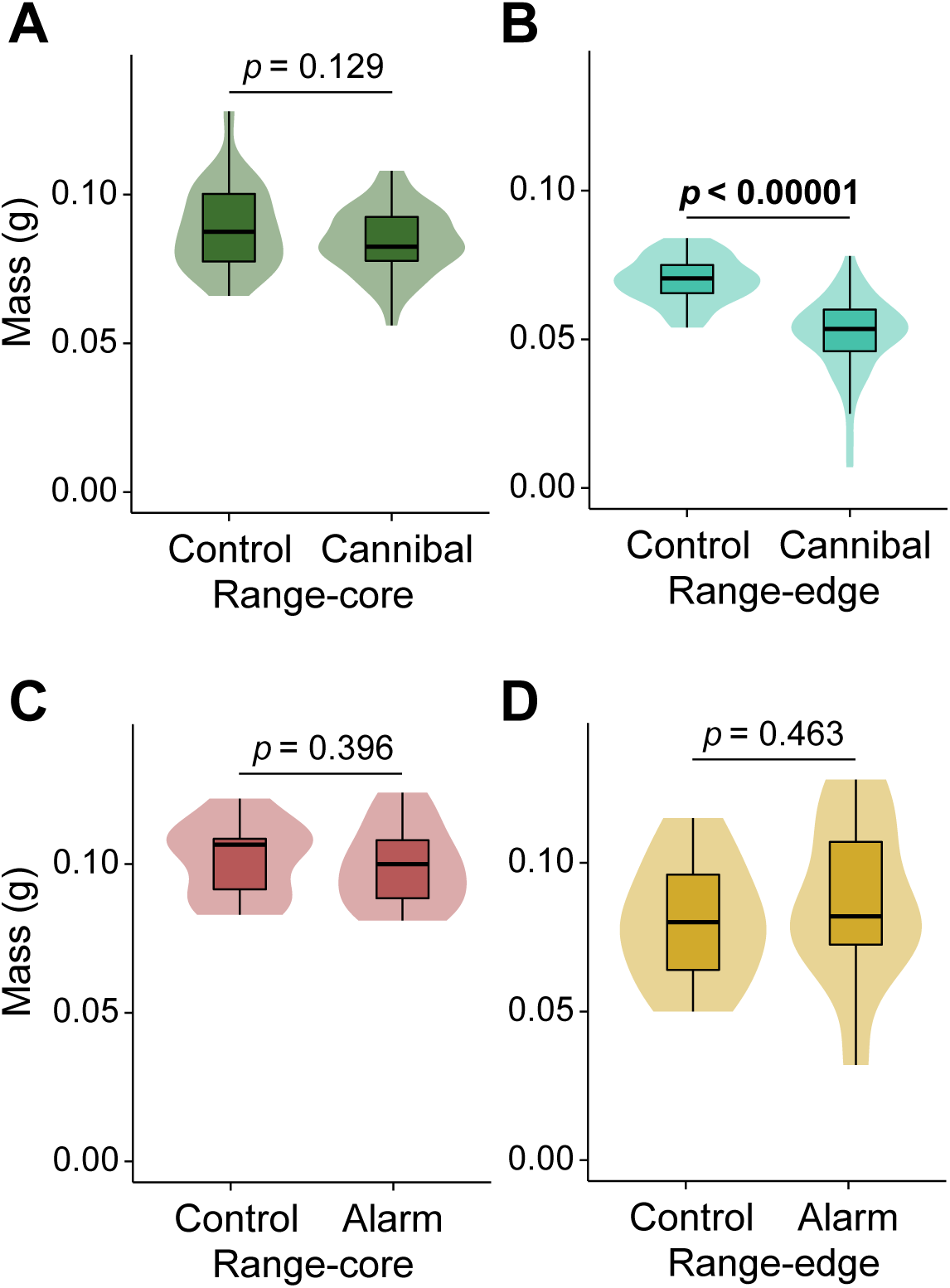
Effect of exposure to conspecific cues on tadpole growth. Mass (g) of tadpoles exposed to cannibal cues and controls from (**A**) range-core and (**B**) range-edge populations, and of tadpoles exposed to alarm cues and controls from (**C**) range-core and (**D**) range-edge populations. Violin plots represent median, interquartile range (IQR), 1.5 × IQR, and kernel density plot. Significant *p*-values (LMMs) are highlighted in bold.

After filtering and quality trimming, the breadth of coverage (i.e., the number of CpGs with a coverage ≥ 10x) was 2.6 ± 0.1 million CpGs (mean ± SE) for tadpoles from the cannibal cue experiment, and 2.5 ± 0.1 million CpGs for tadpoles from the alarm cue experiment (Table S1). The depth of coverage was respectively 16.6 ± 0.1 and 16.8 ± 0.2 fold (Table S1). The methylation density (i.e., the ratio of mCpGs to CpGs across the covered genome) was very high across all samples, as typically observed in vertebrates (Bird, 2002). Nonetheless, tadpoles exposed to either cannibal cues or alarm cues had higher methylation densities compared to controls in both range-core and range-edge populations (generalized linear mixed effects models [GLMMs], all *p* < 0.00001; Figure 2 A–D). Likewise, the proportion of fully methylated sites was higher in tadpoles exposed to either cannibal cues or alarm cues compared to controls in both range-core and range-edge populations (GLMMs, all *p* < 0.00001; Figure S2 A–D). Finally, the methylation level (i.e., the ratio of C to [C + T] reads at each CpG) was also higher in tadpoles exposed to either cannibal cues or alarm cues compared to controls in both range-core and range-edge populations (GLMMs, all *p* < 0.00001; Figure 2 E–H).

**Figure 2.**
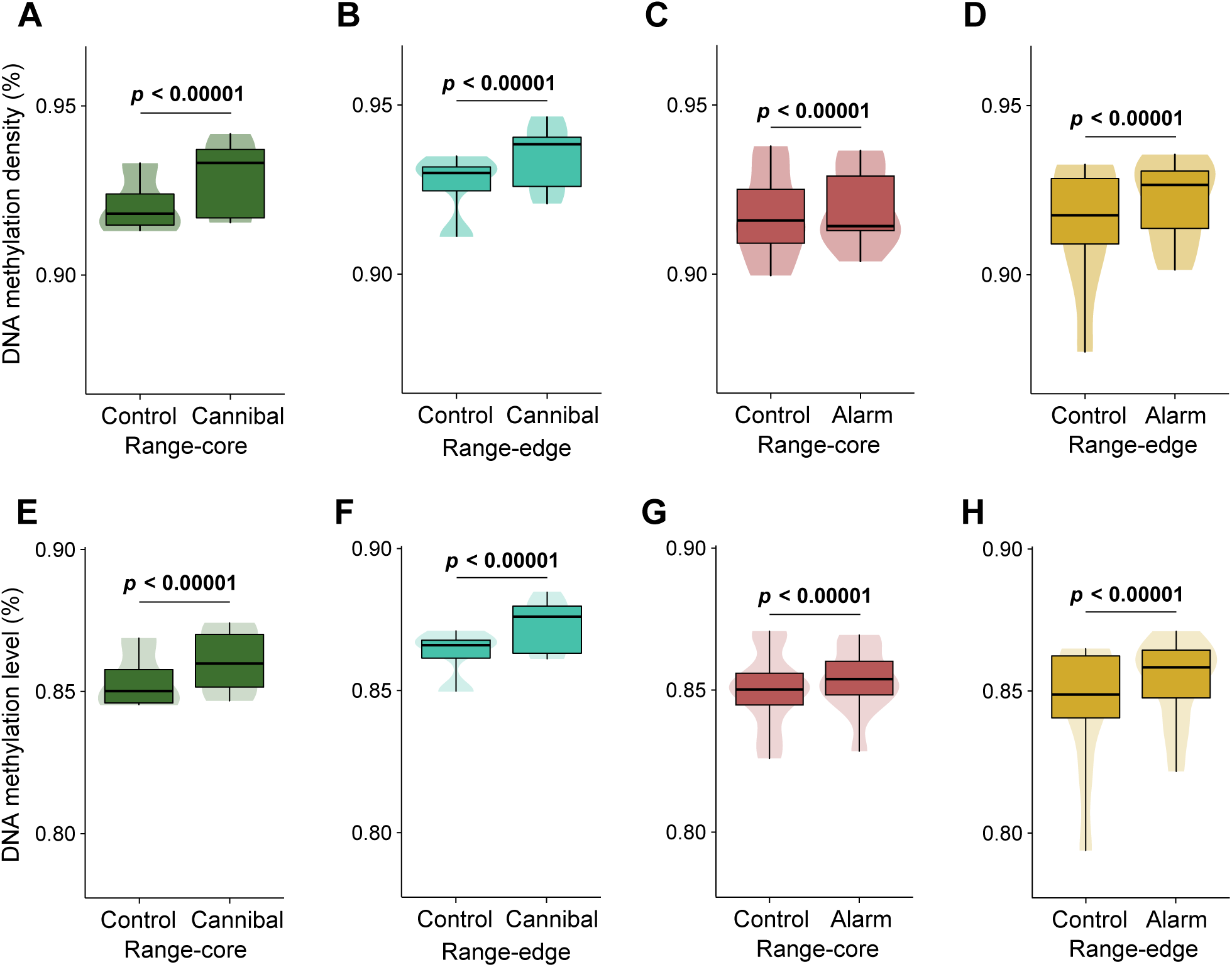
Patterns of DNA methylation in tadpoles exposed to conspecific cues and controls. (**A**–**D**) DNA methylation density (% of mCpGs out of all covered CpGs) in tadpoles exposed to cannibal cues and controls from (**A**) range-core and (**B**) range-edge populations, and in tadpoles exposed to alarm cues and controls from (**C**) range-core and (**D**) range-edge populations. (**E**–**H**) DNA methylation level (% of C out of [C + T] reads at each CpG) in tadpoles exposed to cannibal cues and controls from (**E**) range-core and (**F**) range-edge populations, and in tadpoles exposed to alarm cues and controls from (**G**) range-core and (**H**) range-edge populations. Violin plots represent median, interquartile range (IQR), 1.5 × IQR, and kernel density plot. Significant *p*-values (GLMMs) are highlighted in bold.

Hierarchical clustering based on methylation levels revealed a clear clustering by clutch identity for both range-core and range-edge populations (Figures S3 and S4). This pattern indicates that genetic differences were the main driver of epigenetic differences between samples, whereas treatments had a comparatively minor (albeit statistically significant) effect.

Across all groups, the number of DMCs in cue-exposed tadpoles compared to controls ranged from 13,610 to 50,747, of which 58.4–62.8% were hypermethylated in tadpoles exposed to either cannibal cues or alarm cues. Most DMCs (90.0% and 89.4% of hypermethylated and hypomethylated DMCs, respectively) were specific to tadpoles from one population exposed to one treatment (Figure S5). Only 11 DMCs had higher methylation levels across all cue-exposed tadpoles compared to controls, six of them intersecting with genes *LPAR1-B*, *PSMD10*, *SPC25*, *CASK*, and uncharacterised transcripts *RMA_00056855* and *RMA_00042636* (Figure S5A). Likewise, only four hypomethylated DMCs were common across all groups, two of them intersecting with genes *FMNL2* and *TPX2-B* (Figure S5B).

There were 34 DMRs in range-core tadpoles exposed to cannibal cues compared to controls, of which 20 (58.8%) were located within genes, and 1 (2.9%) was found in the promoter region of an uncharacterised transcript (Table S2). In range-edge tadpoles exposed to cannibal cues, there were 74 DMRs compared to controls, out of which 30 (40.5%) were found within genes, and 6 (8.1%) were located in promoter regions (Table S3). DMRs each contained on average 5.4 ± 2.9 DMCs (mean ± SD; range 3–13) and 5.9 ± 3.0 DMCs (range 3–17) in range-core and range-edge tadpoles, respectively. The majority of DMRs were hypermethylated in cannibal-cue-exposed tadpoles compared to controls in both populations (respectively 64.7% and 60.8%; Tables S2 and S3).

Only 6 DMRs overlapped across both populations in tadpoles exposed to cannibal cues compared to controls (Figure 3), out of which 4 were located within genes *FAM168A*, *RYK*, *MAPK14*, and *HYDIN*. Further, only 2 overlapping DMRs (intersecting with *FAM168A* and *MAPK14*) showed parallel changes in DNA methylation levels in both populations, while the other 4 showed opposite changes in range-core and range-edge tadpoles compared to controls.

**Figure 3.**
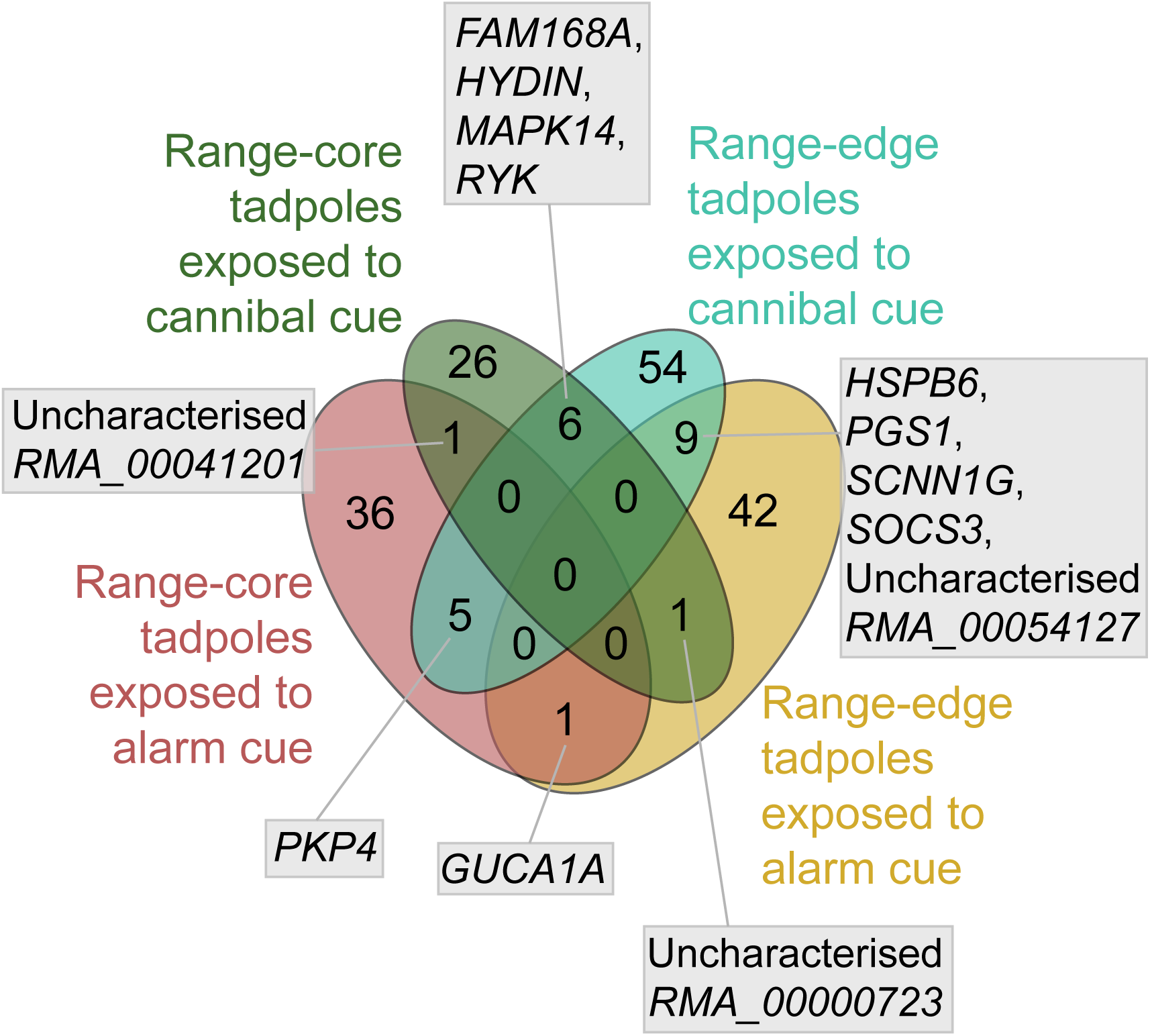
DNA methylation changes in tadpoles exposed to cannibal cues and alarm cues. Venn diagram represents the overlap of DMRs in cannibal-cue-exposed tadpoles versus controls and in alarm-cue-exposed tadpoles versus controls, both from range-core and range-edge populations. Genes intersecting with overlapping DMRs are indicated.

There were 43 DMRs between range-core tadpoles exposed to alarm cues and controls, out of which 19 (44.2%) were found within genic regions, and 5 (11.6%) were found in promoter regions (Table S4). In range-edge tadpoles exposed to alarm cues, there were 53 DMRs compared to controls, out of which 27 (50.9%) were located within genes, and 2 (3.8%) were found in promoter regions (Table S5). Each DMR contained on average 5.1 ± 2.6 DMCs (range 3–15) in range-core tadpoles, and 6.6 ± 4.2 DMCs (range 3–25) in range-edge tadpoles. As for cannibal-cue-exposed tadpoles, most DMRs showed a significant increase in DNA methylation level compared to controls in both range-core and range-edge populations (respectively 72.1% and 66.0%).

Only 1 DMR, located within the *GUCA1A* gene, overlapped and showed an increase in DNA methylation levels across both populations in tadpoles exposed to alarm cues compared to controls (Figure 3).

Overall, there was minimal overlap in regions showing differential methylation in tadpoles exposed to cannibal cues and in tadpoles exposed to alarm cues compared to controls (Figure 3). Only 1 DMR overlapped between cannibal-cue-exposed and alarm-cue-exposed tadpoles from range-core populations. This DMR was located in the promoter region of an uncharacterised transcript, *RMA_00041201*, and showed hypermethylation in both treatments relative to controls. In range-edge tadpoles, 9 DMRs overlapped between tadpoles exposed to cannibal cues and those exposed to alarm cues. Five of those DMRs intersected with genes *HSPB6*, *PGS1*, *SCNN1G*, *SOCS3*, as well as the uncharacterised transcript *RMA_00054127*. However, only genes *PGS1*, *SCNN1G* and *SOCS3* exhibited parallel changes in DNA methylation levels in both treatments relative to controls (hypomethylation for *PGS1* and *SOCS3* and hypermethylation for *SCNN1G*). Of the 4 remaining intergenic DMRs, 2 showed parallel changes and 2 showed opposite changes in DNA methylation levels in both treatments relative to controls.

One DMR (showing opposite changes in DNA methylation levels and intersecting with uncharacterised transcript *RMA_00000723*) overlapped between range-core tadpoles exposed to cannibal cues and range-edge tadpoles exposed to alarm cues. Lastly, 5 DMRs overlapped between range-edge tadpoles exposed to cannibal cues and range-core tadpoles exposed to alarm cues. Only 1 of those DMRs was located within gene *PKP4*. Furthermore, 3 of those DMRs (including *PKP4*) showed opposite changes in DNA methylation levels in both treatments relative to controls.

There was no DMR overlap across both treatments and both populations (Figure 3). GO enrichment analyses did not reveal any significantly over-represented GO term for all DMR lists.

## Discussion

Our findings reveal that exposure to conspecific cannibal cues and alarm cues both changed the DNA methylation profiles of cane toad tadpoles, but did so in different ways. Our results thus suggest that the developmental plastic responses seen in these two contexts, despite their similar short- and long-term consequences, are underpinned by distinct molecular mechanisms.

We did observe a similar overall pattern of hypermethylation in tadpoles exposed to both conspecific cues in both populations compared to controls. These slightly higher levels of DNA methylation might indicate marginally lower levels of gene expression (Jaenisch & Bird, 2003) in tadpoles exposed to conspecific cues. Nonetheless, these changes were small (typically < 1%), and the relationship between DNA methylation levels and transcriptional activity are far from being universal and unidirectional (de Mendoza, Lister, & Bogdanovic, 2020).

We also observed some overlap in DMRs between cannibal-cue-exposed tadpoles compared to controls and alarm-cue-exposed tadpoles compared to controls. One DMR overlapped between both treatments in range-core tadpoles, and 9 DMRs overlapped between both treatments in range-edge tadpoles. These DMRs were found within the gene bodies of *HSPB6*, *PGS1*, *SCNN1G*, *SOCS3*, as well as two uncharacterised genes. *HSPB6* is involved in the regulation of angiogenesis (Vafiadaki, Arvanitis, Eliopoulos, Kranias, & Sanoudou, 2020), *SCNN1G* plays a role in water homeostasis (Hobbs et al., 2013), while *SOCS3* is involved in the regulation of food intake (Zhu et al., 2021). DNA methylation patterns of 5’-flanking regions have been shown to correlate with gene expression, at least for *SCNN1G* (Pierandrei et al., 2021). Changes in DNA methylation in these genes were already evidenced in cane toad tadpoles following exposure to alarm cues (Sarma et al., 2021; Sarma et al., 2020), and we here show that the same modifications occur after exposure to cannibal cues. These findings offer a possible mechanistic link explaining why individuals exposed to alarm cues produce offspring that emit more potent cannibal cues (Sarma et al., 2021).

Changes in DNA methylation levels in the above-mentioned genes may well have developmental consequences during cane toads’ larval life. Yet, as well as being restricted to only a few overlapping regions, changes in DNA methylation levels were not always consistent across treatments (e.g., hypermethylation in one case and hypomethylation in the other). This casts doubts on the causative link between changes in DNA methylation levels, changes in gene expression levels, and downstream phenotypic consequences due to exposure to cannibal cues and alarm cues. It is interesting that some DMCs overlapping across all treatments and populations were found in development-related genes, including *LPAR1-B*, involved in neurogenesis (Fukushima, Kimura, & Chun, 1998), and *FMNL2*, which plays a role in cell morphogenesis (Bai et al., 2011). Nonetheless, it is unclear mechanistically how changes in DNA methylation levels of single CpGs can affect gene activity [but see e.g., Sobiak & Leśniak (2019)].

Within each treatment, additional DMRs intersected with genes also putatively involved in developmental processes. For example, cannibal-cue-exposed tadpoles showed differences in DNA methylation levels in the genes *AKTIP-A*, *HDYN*, *LARGE1*, *LRP4*, *MAPK14*, *NIN*, *NLK*, *PLXNA1, RANBP3L*, *RYK*, *SEZ6* and *SOX5*. *AKTIP-A* mouse mutants show developmental abnormalities (Anselme, Laclef, Lanaud, Rüther, & Schneider-Maunoury, 2007). *HDYN*, *PLXNA1* and SOX5 play a role in brain development (Andrews, Davidson, Tamamaki, Ruhrberg, & Parnavelas, 2016; Palmer et al., 2016; Stolt et al., 2006). *LARGE1*, *MAPK14*, *NLK*, *RANBP3L* and *RYK* are involved in skeletal system development (Chen et al., 2015; Goddeeris et al., 2013; Greenblatt et al., 2010; Halford et al., 2000; Zanotti & Canalis, 2012). *LRP4* plays a role in normal organ development (Weatherbee, Anderson, & Niswander, 2006). *NIN* is integral to epidermis development (Lecland, Hsu, Chemin, Merdes, & Bierkamp, 2019). *SEZ6* is involved in the regulation of motor functions (Gunnersen et al., 2020). There was further hypomethylation in the promoter of *GRM7*, a gene involved in conditioned fear response (Masugi et al., 1999). Alarm-cue-exposed tadpoles showed differences in DNA methylation levels in the genes *CEP135*, *DACH2*, *ECE1*, *EIF3A*, *ENG*, *GPR155*, *MINK1*, *MYL3*, *TIE1* and *ZCCHC3*. *CEP135* and *MINK1* are involved in brain development (Bamborschke et al., 2020; Dan et al., 2000). *DACH2* plays a role in muscle development (Tang & Goldman, 2006). *ECE1*, *ENG* and *MYL3* are involved in heart development (Arthur et al., 2000; James et al., 1999; Poltavski et al., 2019). *EIF3A* plays a role in intestinal development (Liu et al., 2007). *TIE1* is involved in angiogenesis (Loughna & Sato, 2001). *ZCCHC3* plays a role in innate immune response (Lian et al., 2018). Finally, *GPR155* is involved in cognitive functions (Nishimura et al., 2007). Overall, these genes are interesting candidates for future studies, whose main focus should be directed towards generating gene expression data for tadpoles exposed to both conspecific cues and controls across development. This should help to confirm that the above-mentioned genes that show differences in DNA methylation levels between treatments also show differences in gene expression levels, and should bring us one step closer to establishing a causal link between molecular mechanisms and developmental plasticity.

Our results revealed clearly that epigenetic differences between tadpoles were mostly driven by their clutch identity. This phenomenon has previously been documented in cane toads (Sarma et al., 2020), and indicates that genetic differences between individuals are the main cause for their divergence in DNA methylation patterns. The influence of genotypic variation on DNA methylation marks appears ubiquitous. Mounting evidence shows that, although epigenetic marks can be modified by environmental exposure, in many cases they do so under genetic control (Do et al., 2017; Gaunt et al., 2016; Hannon et al., 2018; Kerkel et al., 2008; Min et al., 2021; Tycko, 2010; Villicaña & Bell, 2021). These results stress the importance of controlling for genetic effects (i.e., having a balanced experimental design in terms of clutch identity) when investigating differences in DNA methylation patterns between treatment and control. These results also help explain why we observed only minimal overlap in DMRs between range-core and range-edge tadpoles, even within each treatment (i.e., for tadpoles exposed to the same conspecific cue). Because tadpoles from distinct populations necessarily came from distinct clutches, their constitutive genetic differences induced population-specific DNA methylation patterns (prior to any conspecific cue exposure) that were of greater magnitude than the effect of treatment itself. Our findings further complement previous results showing that clutches vary in their reaction norms, i.e., in their propensity to accelerate their development, when exposed to conspecific cues (DeVore, Crossland, & Shine, 2021).

We found that changes in DNA methylation following exposure to both cannibal cues and alarm cues were largely population-specific, and were of greater magnitude in range-edge tadpoles than in range-core tadpoles. This effect was mirrored in the developmental effects of conspecific cue exposure (i.e., greater effect on mass at the range-edge). These population-specific DNA methylation patterns are consistent with previous studies investigating epigenetic patterns in cane toads (Sarma et al., 2021; Sarma et al., 2020). More generally, they are consistent with the well documented between-population differences in cane toad morphology (Hudson, McCurry, Lundgren, McHenry, & Shine, 2016; Phillips, Brown, Webb, & Shine, 2006), physiology (Brown, Kelehear, Shilton, Phillips, & Shine, 2015), behaviour (Gruber, Brown, Whiting, & Shine, 2017; Lindstrom, Brown, Sisson, Phillips, & Shine, 2013), transcriptomics (Rollins, Richardson, & Shine, 2015; Yagound et al., 2022; Yagound et al., 2022) and genetics (Selechnik et al., 2019) across the Australian invasive range.

We did not detect any significant reduction in body mass in alarm-cue-exposed tadpoles. By contrast, previous studies found such an effect at a later stage in development (i.e., at metamorphosis) (Crossland et al., 2019; Hagman et al., 2009; Hagman & Shine, 2009). Thus, the apparent lack of growth reduction seen following alarm-cue exposure may be an artefact of early euthanasia.

Exposure to cannibal cues and alarm cues thus appear to trigger distinct molecular mechanisms. Both cues affect DNA methylation patterns locally, but each in largely distinct genomic regions. It is interesting to contrast these findings with the observations that both cues trigger reduced growth responses in tadpoles (Crossland & Shine, 2012; DeVore, Crossland, & Shine, 2021; Hagman et al., 2009; Hagman & Shine, 2009), and that hatchlings exposed to alarm cues have offspring that themselves produce more potent cannibal cues (Sarma et al., 2021). Several hypotheses might explain this discrepancy. Each cue might trigger a series of molecular changes involving many genes within complex networks. It is possible that each cue does indeed involve distinct causal molecular mechanisms that result in similar phenotypic effects. While growth reduction in tadpoles is a direct consequence in the case of exposure to alarm cues (Crossland et al., 2019; Hagman et al., 2009; Hagman & Shine, 2009), it is a carry-over effect of the hatchling stage in the case of exposure to cannibal cues (Clarke et al., 2015; Crossland & Shine, 2012; DeVore, Crossland, & Shine, 2021), which might contribute to the lack of overlap in DNA methylation changes seen between exposure to both conspecific cues. By contrast, it is also possible that these gene networks are quite similar between both contexts, but that we were only able to capture a fraction of the genomic regions involved in each case. The lack of overlap could then derive from constraints in our experimental design in terms of sample size, statistical power, sequencing methodology, breadth of coverage, and/or underlying genetic differences. If the molecular changes underlying developmental plasticity are restricted to a short time-window, it is possible that we sampled tadpoles too late to detect them. Our experiments were conducted at different times, and involved tadpoles of slightly different ages, which could also have introduced artefacts in our results. Lastly, is it also possible that the changes we observed in DNA methylation patterns are not causally involved in cane toad developmental plasticity. DNA methylation marks might well be affected by exposure to conspecific cues, but perhaps these changes are by-products of conspecific-cue exposure, or a consequence of other molecular changes (perhaps also epigenetic in nature, such as histone post-translational modifications [Cedar & Bergman, 2009]) that are themselves the cause of downstream developmental changes. Gene expression data matched to epigenetic data are needed to solve this enduring puzzle.

## Acknowledgements

We thank Jack Reid and Simon Ducatez for assistance with experimental work. This work was supported by a Deakin University Fellowship and a UNSW Scientia Fellowship to LAR, and the Australian Research Council (DE150101393 to LAR, FL120100074 to RS, and DP160102991 to RS and LAR).

## Data Accessibility and Benefit-Sharing Statement

The RRBS data for the 26 samples from the cannibal cue experiment, and the 41 samples from the alarm cue experiment is available at the National Center for Biotechnology Information (NCBI) Sequence Read Archive (BioProject PRJNA901184).

## Author Contributions

MFR, CMRL, JLD, RS and LAR designed research. RRS, MRC and GPB performed research. BY, RRB, RJE and GPB analyzed data. CMRL, MRC, JLD, RS, MRC and LAR contributed to interpretation and writing. BY wrote the paper.

