## Supplementary material for "Is developmental plasticity triggered by DNA methylation changes in the invasive cane toad (*Rhinella marina*)?": Figure S1

### Table of Contents:

| | |
|---|---|
| **Figure S1** | Page 2 |
| **Figure S2** | Page 3 |
| **Figure S3** | Page 4 |
| **Figure S4** | Page 5 |
| **Figure S5** | Page 6 |

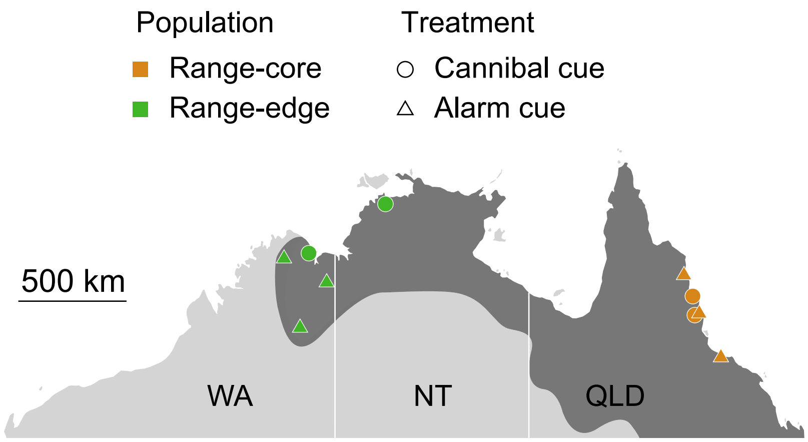


**Figure S1. Location of samples.** Samples used in the cannibal cue experiment originated from localities depicted by circles. Samples used in the alarm cue experiment originated from localities depicted by triangles. Orange and green localities respectively correspond to range-core and range-edge populations. NT, Northern Territory; QLD, Queensland; WA, Western Australia. The shaded area represents the cane toad's Australian invasive range.

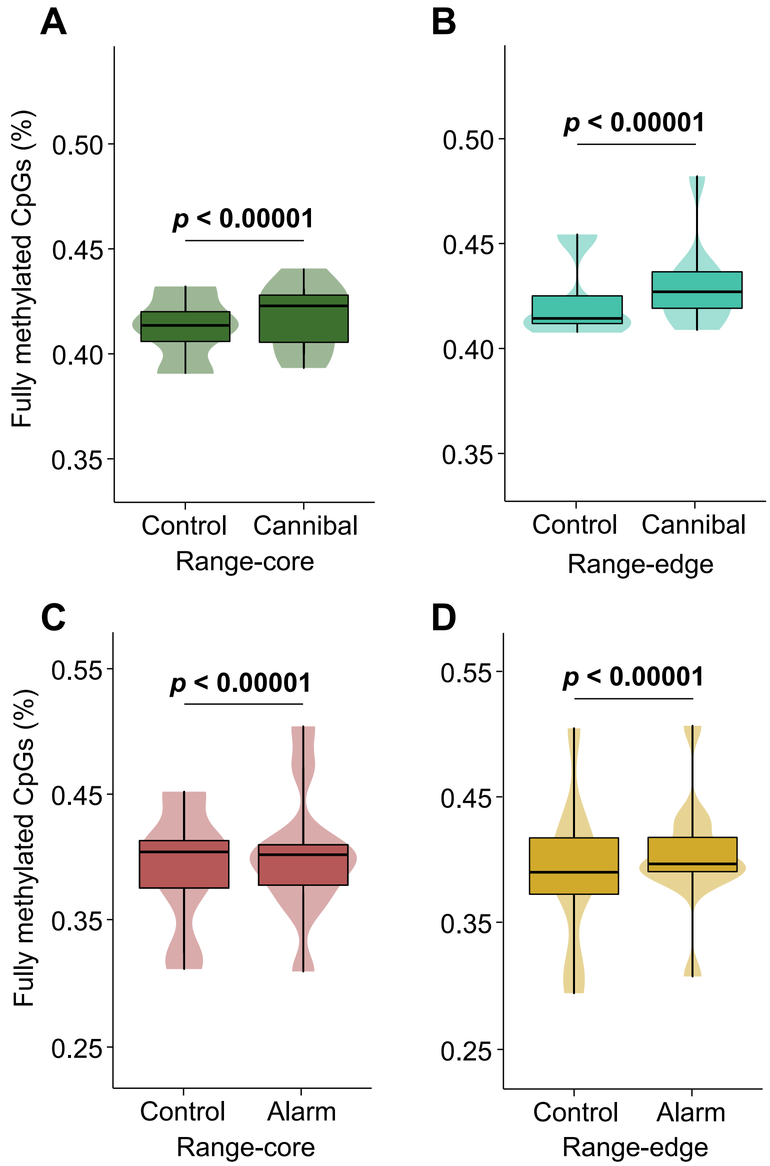


**Figure S2. Patterns of DNA methylation in tadpoles exposed to conspecific cues and controls.** (**A**,**B**) Proportion of fully methylated CpGs (%) in tadpoles exposed to cannibal cues and controls from (**A**) range-core and (**B**) range-edge populations. (**C**,**D**) Proportion of fully methylated CpGs (%) in tadpoles exposed to alarm cues and controls from (**C**) range-core and (**D**) range-edge populations. Violin plots represent median, interquartile range (IQR), 1.5 × IQR, and kernel density plot. Significant *p*-values (GLMMs) are highlighted in bold.

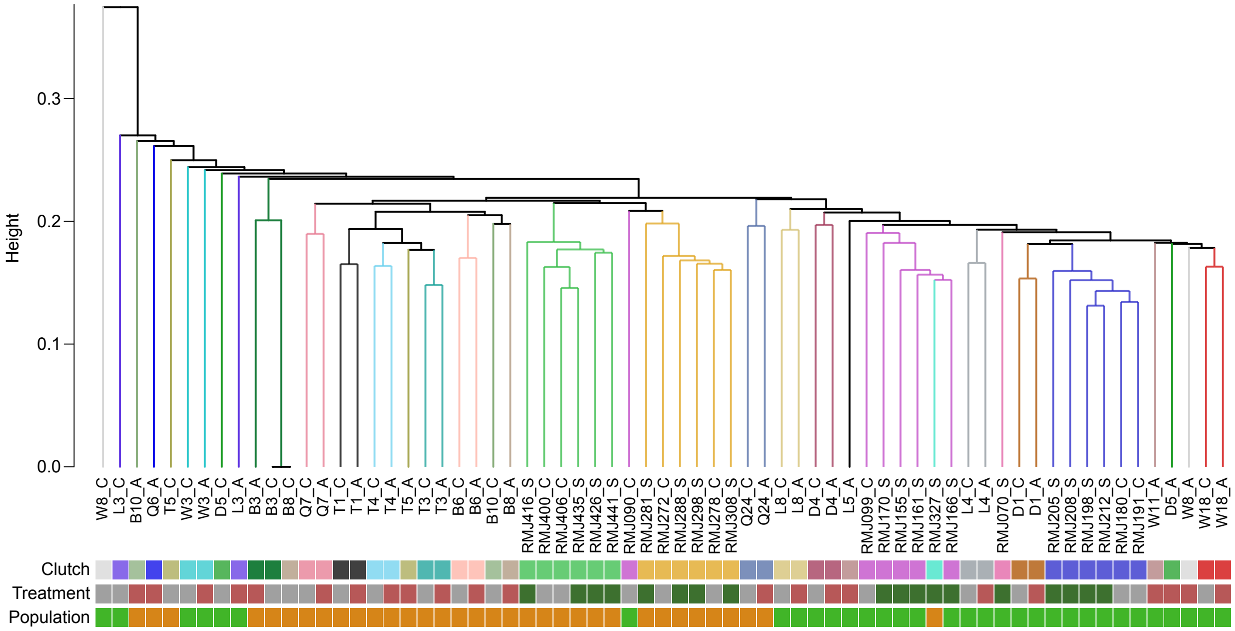


**Figure S3. Hierarchical clustering (average agglomerative method on correlation distances) of DNA methylation status for all samples.** Coloured boxes group samples by clutch, treatment (i.e., red, alarm cue; dark green, cannibal cue; grey, control) and population (i.e., orange, range-core; light green, range-edge).

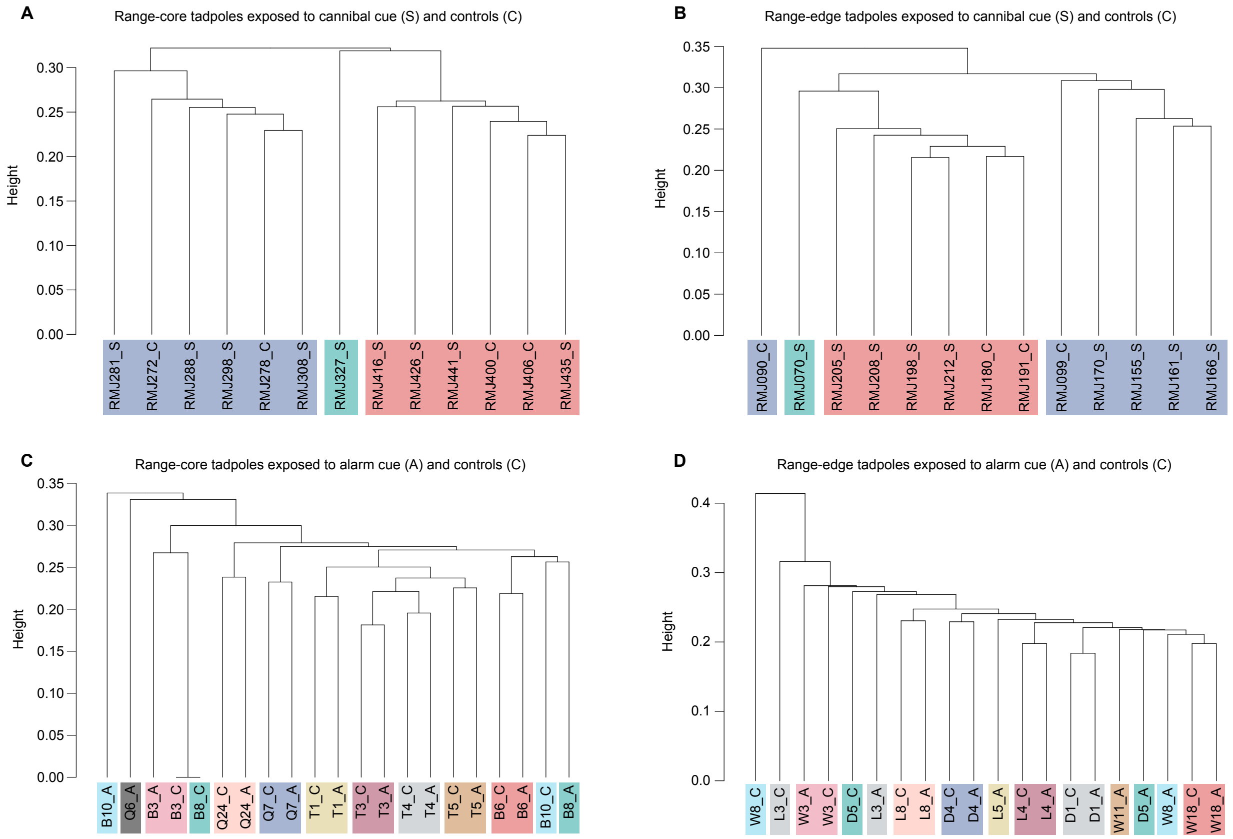


**Figure S4. Clustering of samples according to their DNA methylation status.** (**A**–**D**) Hierarchical clustering (average agglomerative method on correlation distances) of DNA methylation status for (**A**) range-core and (**B**) range-edge tadpoles exposed to cannibal cues and controls, and (**C**) range-core and (**D**) range-edge tadpoles exposed to alarm cues and controls. Different colours represent different clutches.

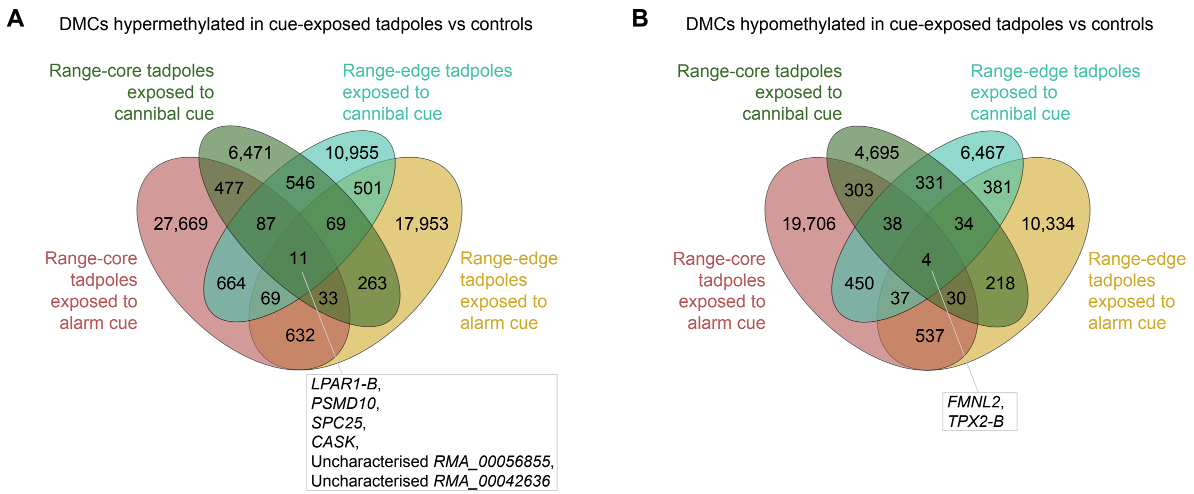


**Figure S5. Local DNA methylation changes in tadpoles exposed to cannibal cues and alarm cues.** Venn diagram represent the overlap of (**A**) hypermethylated and (**B**) hypomethylated DMCs in cannibal-cue-exposed tadpoles versus controls and in alarm-cue-exposed tadpoles versus controls, both from range-core and range-edge populations. Genes intersecting with overlapping DMCs across all groups are indicated.
