## Supplementary material for "Is developmental plasticity triggered by DNA methylation changes in the invasive cane toad (*Rhinella marina*)?": Table S1

**Table S1.** DNA methylation statistics for each sample.

| Individual | Population | Treatment | Coverage (mean ± SE) | No. covered CpGs (> 10x) | No. fully methylated CpGs (%) | DNA methylation level (%, mean ± SE) |
| --- | --- | --- | --- | --- | --- | --- |
| B10_C | Range-core | Control | 17.11 ± 0.01 | 2,706,712 | 1,012,241 (37.40) | 85.19 ± 0.01 |
| B10_A | Range-core | Alarm cue | 15.82 ± 0.01 | 1,851,760 | 870,936 (47.03) | 84.49 ± 0.02 |
| B3_C | Range-core | Control | 16.07 ± 0.01 | 2,120,158 | 956,668 (45.12) | 84.52 ± 0.02 |
| B3_A | Range-core | Alarm cue | 17.63 ± 0.01 | 2,882,616 | 1,198,234 (41.57) | 86.43 ± 0.01 |
| B6_C | Range-core | Control | 17.08 ± 0.01 | 2,685,849 | 1,099,764 (40.95) | 86.34 ± 0.01 |
| B6_A | Range-core | Alarm cue | 19.01 ± 0.01 | 3,466,836 | 1,396,244 (40.27) | 86.30 ± 0.01 |
| B8_C | Range-core | Control | 16.07 ± 0.01 | 2,120,158 | 956,668 (45.12) | 84.52 ± 0.02 |
| B8_A | Range-core | Alarm cue | 16.45 ± 0.01 | 2,597,621 | 996,799 (38.37) | 84.89 ± 0.02 |
| Q24_C | Range-core | Control | 16.27 ± 0.01 | 2,277,887 | 943,133 (41.40) | 84.47 ± 0.02 |
| Q24_A | Range-core | Alarm cue | 17.21 ± 0.01 | 2,377,544 | 955,539 (40.19) | 85.90 ± 0.01 |
| Q6_A | Range-core | Alarm cue | 16.01 ± 0.01 | 1,897,238 | 952,975 (50.23) | 84.99 ± 0.02 |
| Q7_C | Range-core | Control | 15.66 ± 0.01 | 2,062,043 | 834,106 (40.45) | 83.82 ± 0.02 |
| Q7_A | Range-core | Alarm cue | 16.30 ± 0.01 | 2,500,833 | 1,009,438 (40.36) | 84.44 ± 0.02 |
| T1_C | Range-core | Control | 18.49 ± 0.01 | 2,854,455 | 1,152,233 (40.37) | 86.57 ± 0.01 |
| T1_A | Range-core | Alarm cue | 16.82 ± 0.01 | 2,911,986 | 1,139,738 (39.14) | 85.13 ± 0.01 |
| T3_C | Range-core | Control | 17.07 ± 0.01 | 2,704,868 | 845,071 (31.24) | 82.12 ± 0.02 |
| T3_A | Range-core | Alarm cue | 16.99 ± 0.01 | 3,025,076 | 1,087,944 (35.96) | 83.83 ± 0.02 |
| T4_C | Range-core | Control | 16.91 ± 0.01 | 3,188,903 | 1,216,855 (38.16) | 84.82 ± 0.01 |
| T4_A | Range-core | Alarm cue | 16.94 ± 0.01 | 2,712,431 | 842,394 (31.06) | 82.37 ± 0.02 |
| T5_C | Range-core | Control | 17.95 ± 0.01 | 2,149,651 | 699,234 (32.53) | 82.87 ± 0.02 |
| T5_A | Range-core | Alarm cue | 16.04 ± 0.01 | 2,820,292 | 1,050,226 (37.24) | 84.23 ± 0.02 |
| D1_C | Range-edge | Control | 17.76 ± 0.01 | 3,379,797 | 1,255,520 (37.15) | 85.42 ± 0.01 |
| D1_A | Range-edge | Alarm cue | 18.33 ± 0.01 | 3,503,050 | 1,367,471 (39.04) | 86.13 ± 0.01 |
| D4_C | Range-edge | Control | 17.46 ± 0.01 | 2,711,875 | 1,126,971 (41.56) | 86.28 ± 0.01 |
| D4_A | Range-edge | Alarm cue | 16.46 ± 0.01 | 2,493,594 | 765,879 (30.71) | 81.99 ± 0.02 |
| D5_C | Range-edge | Control | 15.31 ± 0.01 | 1,606,267 | 472,261 (29.40) | 79.22 ± 0.02 |
| D5_A | Range-edge | Alarm cue | 16.84 ± 0.01 | 2,979,308 | 1,155,748 (38.79) | 85.62 ± 0.01 |
| L3_C | Range-edge | Control | 16.13 ± 0.01 | 1,772,721 | 888,587 (50.13) | 84.67 ± 0.02 |
| L3_A | Range-edge | Alarm cue | 15.29 ± 0.01 | 1,688,226 | 668,014 (39.57) | 83.29 ± 0.02 |
| L4_C | Range-edge | Control | 16.52 ± 0.01 | 2,408,495 | 759,380 (31.53) | 81.88 ± 0.02 |
| L4_A | Range-edge | Alarm cue | 16.93 ± 0.01 | 3,183,418 | 1,242,975 (39.05) | 85.67 ± 0.01 |
| L5_A | Range-edge | Alarm cue | 17.68 ± 0.01 | 2,717,554 | 1,104,151 (40.63) | 86.31 ± 0.01 |
| L8_C | Range-edge | Control | 15.65 ± 0.01 | 2,213,397 | 860,310 (38.87) | 83.86 ± 0.02 |
| L8_A | Range-edge | Alarm cue | 15.39 ± 0.01 | 2,035,356 | 804,367 (39.52) | 83.95 ± 0.02 |
| W11_A | Range-edge | Alarm cue | 16.63 ± 0.01 | 2,561,577 | 1,105,457 (43.16) | 86.36 ± 0.01 |
| W18_C | Range-edge | Control | 17.49 ± 0.01 | 3,559,635 | 1,380,915 (38.79) | 86.02 ± 0.01 |
| W18_A | Range-edge | Alarm cue | 17.22 ± 0.01 | 2,638,045 | 1,122,813 (42.56) | 86.88 ± 0.01 |
| W3_C | Range-edge | Control | 15.61 ± 0.01 | 1,268,902 | 547,030 (43.11) | 86.14 ± 0.02 |
| W3_A | Range-edge | Alarm cue | 16.75 ± 0.01 | 2,395,814 | 1,205,656 (50.32) | 85.37 ± 0.02 |
| W8_C | Range-edge | Control | 18.27 ± 0.03 | 676,604 | 268,837 (39.73) | 84.33 ± 0.03 |
| W8_A | Range-edge | Alarm cue | 16.50 ± 0.01 | 3,059,678 | 1,175,334 (38.41) | 85.16 ± 0.01 |
| RMJ272_C | Range-core | Control | 15.41 ± 0.01 | 2,023,970 | 840,345 (41.52) | 84.89 ± 0.02 |
| RMJ278_C | Range-core | Control | 16.07 ± 0.01 | 2,578,716 | 1,057,554 (41.01) | 85.67 ± 0.02 |
| RMJ281_S | Range-core | Cannibal cue | 16.30 ± 0.01 | 2,114,801 | 930,636 (44.01) | 85.07 ± 0.02 |
| RMJ288_S | Range-core | Cannibal cue | 15.93 ± 0.01 | 2,377,902 | 948,244 (39.88) | 84.93 ± 0.02 |
| RMJ298_S | Range-core | Cannibal cue | 15.84 ± 0.01 | 2,246,608 | 948,199 (42.21) | 85.42 ± 0.02 |
| RMJ308_S | Range-core | Cannibal cue | 17.11 ± 0.01 | 2,761,054 | 1,170,417 (42.39) | 87.17 ± 0.01 |
| RMJ327_S | Range-core | Cannibal cue | 17.24 ± 0.01 | 3,267,404 | 1,405,491 (43.02) | 87.68 ± 0.01 |
| RMJ400_C | Range-core | Control | 16.95 ± 0.01 | 2,696,471 | 1,163,267 (43.14) | 87.14 ± 0.01 |
| RMJ406_C | Range-core | Control | 16.29 ± 0.01 | 2,641,992 | 1,028,735 (38.94) | 84.79 ± 0.02 |
| RMJ416_S | Range-core | Cannibal cue | 17.40 ± 0.01 | 2,280,409 | 974,285 (42.72) | 87.50 ± 0.01 |
| RMJ426_S | Range-core | Cannibal cue | 17.44 ± 0.01 | 2,760,301 | 1,082,488 (39.22) | 86.25 ± 0.01 |
| RMJ435_S | Range-core | Cannibal cue | 16.48 ± 0.01 | 2,454,708 | 993,000 (40.45) | 85.69 ± 0.02 |
| RMJ441_S | Range-core | Cannibal cue | 18.35 ± 0.01 | 2,884,432 | 1,192,090 (41.33) | 87.27 ± 0.01 |
| RMJ070_S | Range-edge | Cannibal cue | 16.96 ± 0.01 | 2,452,473 | 1,067,309 (43.52) | 88.06 ± 0.01 |
| RMJ090_C | Range-edge | Control | 15.72 ± 0.01 | 2,365,636 | 975,369 (41.23) | 84.59 ± 0.02 |
| RMJ099_C | Range-edge | Control | 16.19 ± 0.01 | 2,410,288 | 1,088,735 (45.17) | 86.14 ± 0.02 |
| RMJ155_S | Range-edge | Cannibal cue | 16.26 ± 0.01 | 2,449,011 | 1,062,206 (43.37) | 85.90 ± 0.02 |
| RMJ161_S | Range-edge | Cannibal cue | 16.08 ± 0.01 | 2,714,595 | 1,101,465 (40.58) | 85.73 ± 0.01 |
| RMJ166_S | Range-edge | Cannibal cue | 16.28 ± 0.01 | 2,898,435 | 1,206,304 (41.62) | 85.92 ± 0.01 |
| RMJ170_S | Range-edge | Cannibal cue | 16.61 ± 0.01 | 2,583,042 | 1,239,096 (47.97) | 86.31 ± 0.01 |
| RMJ180_C | Range-edge | Control | 16.38 ± 0.01 | 2,853,992 | 1,170,950 (41.03) | 86.70 ± 0.01 |
| RMJ191_C | Range-edge | Control | 16.45 ± 0.01 | 3,063,958 | 1,239,234 (40.45) | 86.27 ± 0.01 |
| RMJ198_S | Range-edge | Cannibal cue | 17.37 ± 0.01 | 3,081,851 | 1,284,695 (41.69) | 87.20 ± 0.01 |
| RMJ205_S | Range-edge | Cannibal cue | 17.52 ± 0.01 | 2,671,839 | 1,093,008 (40.91) | 87.62 ± 0.01 |
| RMJ208_S | Range-edge | Cannibal cue | 16.54 ± 0.01 | 2,653,964 | 1,125,590 (42.41) | 87.20 ± 0.01 |
| RMJ212_S | Range-edge | Cannibal cue | 17.38 ± 0.01 | 2,990,400 | 1,270,081 (42.47) | 87.57 ± 0.01 |

**Table S2.** DMRs between range-core tadpoles exposed to cannibal cues and controls.

| Contig | Start DMR | End DMR | Length DMR (bp) | Methylation difference (%) | No. CpGs | No. DMCs | DMR  *q*-value | Gene | Protein |
| --- | --- | --- | --- | --- | --- | --- | --- | --- | --- |
| ctg7929 | 70,227 | 70,401 | 175 | -41.55 | 9 | 8 | < 0.00001 | *RBKS* | Ribokinase |
| ctg399**^b^** | 944,763 | 945,066 | 304 | -36.11 | 15 | 11 | < 0.00001 | *RMA_00000723* | Unknown |
| ctg6739 | 25,009 | 25,071 | 63 | -33.86 | 6 | 6 | < 0.00001 | Intergenic | N/A |
| ctg3195 | 95,407 | 95,827 | 421 | -28.54 | 15 | 8 | < 0.00001 | *AKTIP-A* | AKT-interacting protein homolog A |
| ctg11181 | 114,196 | 114,447 | 252 | -27.27 | 10 | 8 | < 0.00001 | Intergenic | N/A |
| ctg3933 | 41,095 | 41,188 | 94 | -24.24 | 8 | 4 | 0.00001 | *TXN* | Thioredoxin |
| ctg14726**^c^** | 68,308 | 68,398 | 91 | -24.17 | 6 | 3 | 0.00013 | *RYK* | Tyrosine-protein kinase RYK |
| ctg15338 | 47,218 | 47,513 | 296 | -23.37 | 13 | 13 | < 0.00001 | *SLC47A1* | Multidrug and toxin extrusion protein 1 |
| ctg3167**^c^** | 901,657 | 902,160 | 504 | -23.06 | 12 | 5 | < 0.00001 | Intergenic | N/A |
| ctg23866 | 3,072 | 3,162 | 91 | -20.59 | 8 | 3 | 0.00013 | Intergenic | N/A |
| ctg4387 | 339,637 | 339,733 | 97 | -20.16 | 6 | 3 | < 0.00001 | Intergenic | N/A |
| ctg12090 | 137,482 | 137,780 | 299 | -20.01 | 7 | 4 | < 0.00001 | *SEZ6* | Seizure protein 6 homolog |
| ctg770 | 188,230 | 188,323 | 94 | 20.05 | 8 | 3 | 0.00021 | *PDE1A* | Calcium/calmodulin-dependent 3',5'-cyclic nucleotide phosphodiesterase 1A |
| ctg998**^c^** | 34,962 | 35,753 | 792 | 20.07 | 19 | 3 | 0.00033 | *HYDIN* | Hydrocephalus-inducing protein homolog |
| ctg21927 | 490 | 537 | 48 | 20.69 | 9 | 4 | 0.0023 | Intergenic | N/A |
| ctg4046 | 135,831 | 136,086 | 256 | 21.93 | 14 | 4 | 0.00002 | Intergenic | N/A |
| ctg3168 | 255,027 | 255,218 | 192 | 24.09 | 8 | 4 | 0.0022 | *SHANK3* | SH3 and multiple ankyrin repeat domains protein 3 |
| ctg76 | 446,909 | 447,163 | 255 | 25.04 | 12 | 4 | < 0.00001 | *TMEM163* | Transmembrane protein 163 |
| ctg3843 | 40,389 | 40,436 | 48 | 26.77 | 5 | 3 | < 0.00001 | Intergenic | N/A |
| ctg29953 | 24,973 | 25,187 | 215 | 26.82 | 8 | 6 | < 0.00001 | Intergenic | N/A |
| ctg143 | 70,476 | 70,895 | 420 | 26.94 | 16 | 10 | < 0.00001 | *RMA_00025525* | Unknown |
| ctg7863**^c^** | 136,579 | 136,654 | 76 | 26.97 | 5 | 3 | < 0.00001 | Intergenic | N/A |
| ctg4646 | 112,284 | 112,345 | 62 | 27.46 | 5 | 3 | 0.00046 | *NIN* | Ninein |
| ctg1638**^c^** | 23,284 | 23,587 | 304 | 28.42 | 9 | 7 | < 0.00001 | *MAPK14* | Mitogen-activated protein kinase 14 |
| ctg25158 | 53,008 | 53,098 | 91 | 28.58 | 5 | 3 | 0.00026 | Intergenic | N/A |
| ctg6921 | 20,708 | 20,779 | 72 | 30.34 | 7 | 6 | < 0.00001 | *SMIM4* | Small integral membrane protein 4 |
| ctg11248 | 90,859 | 90,933 | 75 | 30.51 | 6 | 4 | < 0.00001 | Intergenic | N/A |
| ctg29669 | 10,257 | 10,358 | 102 | 30.89 | 7 | 5 | < 0.00001 | Intergenic | N/A |
| ctg9969**^a^** | 28,668 | 28,807 | 140 | 32.70 | 5 | 4 | < 0.00001 | Promoter of *RMA_00041201* | Unknown |
| ctg3847 | 231,861 | 231,939 | 79 | 33.17 | 5 | 3 | < 0.00001 | *VSTM2B* | V-set and transmembrane domain-containing protein 2B |
| ctg816 | 330,047 | 330,389 | 343 | 34.19 | 13 | 11 | < 0.00001 | *RMA_00012239* | Unknown |
| ctg865 | 445,930 | 446,083 | 154 | 34.83 | 5 | 3 | 0.0012 | *RMA_00005388* | Unknown |
| ctg7585**^c^** | 41,611 | 41,764 | 154 | 37.45 | 12 | 11 | < 0.00001 | *FAM168A* | Protein FAM168A |
| ctg4558 | 49,492 | 49,577 | 86 | 50.27 | 5 | 5 | < 0.00001 | *ADAMTS14* | A disintegrin and metalloproteinase with thrombospondin motifs 14 |

**^a^**Also found between range-core tadpoles exposed to alarm cues and controls.

**^b^**Also found between range-edge tadpoles exposed to alarm cues and controls.

**^c^**Also found between range-edge tadpoles exposed to cannibal cues and controls.

**Table S3.** DMRs between range-edge tadpoles exposed to cannibal cues and controls.

| Contig | Start DMR | End DMR | Length DMR (bp) | Methylation difference (%) | No. CpGs | No. DMCs | DMR  *q*-value | Gene | Protein |
| --- | --- | --- | --- | --- | --- | --- | --- | --- | --- |
| ctg1071 | 247,181 | 247,373 | 193 | -35.69 | 18 | 17 | < 0.00001 | *SOX5* | Transcription factor SOX-5 |
| ctg1072 | 44,611 | 44,778 | 168 | 31.24 | 7 | 4 | < 0.00001 | *RANBP3L* | Ran-binding protein 3-like |
| ctg1164 | 140,743 | 140,808 | 66 | -22.74 | 8 | 4 | 0.0011 | Intergenic | N/A |
| ctg1181**^b^** | 2,961 | 3,336 | 376 | -33.97 | 15 | 9 | < 0.00001 | Intergenic | N/A |
| ctg12002 | 173,499 | 173,586 | 88 | -30.54 | 8 | 8 | < 0.00001 | *RMA_00019178* | Unknown |
| ctg12558 | 33,641 | 34,312 | 672 | 21.03 | 15 | 7 | < 0.00001 | *APEH* | Acylamino-acid-releasing enzyme |
| ctg12829 | 58,878 | 58,963 | 86 | 32.94 | 7 | 7 | < 0.00001 | Promoter of *RMA_00034135* | Unknown |
| ctg13603 | 25,150 | 25,214 | 65 | 29.89 | 6 | 6 | < 0.00001 | Intergenic | N/A |
| ctg13974 | 11,408 | 11,564 | 157 | -21.75 | 8 | 6 | < 0.00001 | Promoter of *GLRA2* | Glycine receptor subunit alpha-2 |
| ctg14020 | 10,370 | 10,474 | 105 | -30.25 | 7 | 6 | < 0.00001 | Promoter of *GRM7* | Metabotropic glutamate receptor 7 |
| ctg14606 | 9,914 | 10,042 | 129 | -28.66 | 7 | 5 | < 0.00001 | Intergenic | N/A |
| ctg14726**^c^** | 68,334 | 68,398 | 65 | 25.06 | 5 | 4 | < 0.00001 | *RYK* | Tyrosine-protein kinase RYK |
| ctg14814 | 121,525 | 121,602 | 78 | 32.10 | 5 | 3 | < 0.00001 | Intergenic | N/A |
| ctg15409 | 4,923 | 5,040 | 118 | 20.13 | 7 | 6 | < 0.00001 | Promoter of *RMA_00024538* | Unknown |
| ctg1590 | 79,976 | 80,019 | 44 | 43.91 | 5 | 5 | < 0.00001 | *LARGE1* | LARGE xylosyl- and glucuronyltransferase 1 |
| ctg16265 | 35,720 | 35,795 | 76 | 34.76 | 6 | 5 | < 0.00001 | *LRP4* | Low-density lipoprotein receptor-related protein 4 |
| ctg1638**^c^** | 23,227 | 23,361 | 135 | 32.19 | 7 | 6 | < 0.00001 | *MAPK14* | Mitogen-activated protein kinase 14 |
| ctg1846 | 86,117 | 86,265 | 149 | -25.13 | 5 | 3 | < 0.00001 | *FOLH1* | Glutamate carboxypeptidase 2 |
| ctg189 | 6,300 | 6,432 | 133 | 22.69 | 8 | 3 | 0.00023 | Intergenic | N/A |
| ctg19321**^b^** | 2,398 | 2,462 | 65 | -22.11 | 6 | 3 | 0.00040 | *RMA_00054127* | Unknown |
| ctg19786**^a^** | 63,399 | 63,623 | 225 | 22.61 | 12 | 5 | 0.00001 | Intergenic | N/A |
| ctg20065 | 18,387 | 18,461 | 75 | 25.88 | 5 | 3 | < 0.00001 | Intergenic | N/A |
| ctg2037 | 59,449 | 59,536 | 88 | -36.60 | 9 | 7 | < 0.00001 | Intergenic | N/A |
| ctg2041 | 250,592 | 250,698 | 107 | 28.26 | 6 | 4 | 0.00009 | Intergenic | N/A |
| ctg21629 | 33,580 | 33,848 | 269 | 27.94 | 6 | 4 | < 0.00001 | Intergenic | N/A |
| ctg2173 | 117,678 | 117,812 | 135 | -30.16 | 8 | 5 | < 0.00001 | *KNL1* | Kinetochore scaffold 1 |
| ctg22578 | 73,170 | 73,239 | 70 | 21.12 | 5 | 3 | 0.0035 | *RMA_00027688* | Unknown |
| ctg22943 | 7,589 | 8,105 | 517 | -22.20 | 16 | 7 | < 0.00001 | Promoter of *RMA_00054644* | Unknown |
| ctg2436 | 4,131 | 4,196 | 66 | -32.54 | 5 | 4 | < 0.00001 | *L1RE1* | LINE-1 retrotransposable element ORF2 protein |
| ctg25051 | 4,062 | 4,376 | 315 | 29.09 | 21 | 14 | < 0.00001 | Intergenic | N/A |
| ctg25970**^b^** | 30,960 | 31,112 | 153 | 39.54 | 13 | 13 | < 0.00001 | Intergenic | N/A |
| ctg26570 | 11,352 | 11,400 | 49 | -29.03 | 5 | 4 | < 0.00001 | Intergenic | N/A |
| ctg2710 | 697,395 | 698,125 | 731 | -23.09 | 28 | 11 | < 0.00001 | *RPTOR* | Regulatory-associated protein of mTOR |
| ctg2897 | 819,247 | 819,342 | 96 | 44.75 | 7 | 7 | < 0.00001 | Intergenic | N/A |
| ctg29626 | 14,134 | 14,606 | 473 | 22.17 | 12 | 10 | < 0.00001 | Intergenic | N/A |
| ctg3167**^c^** | 901,657 | 902,160 | 504 | 31.69 | 12 | 7 | < 0.00001 | Intergenic | N/A |
| ctg3234 | 10,302 | 10,388 | 87 | 31.86 | 8 | 3 | < 0.00001 | Intergenic | N/A |
| ctg3326 | 266,723 | 266,774 | 52 | 26.53 | 6 | 3 | 0.00010 | *PLXNA1* | Plexin-A1 |
| ctg3451 | 4,391 | 4,486 | 96 | 39.43 | 5 | 4 | < 0.00001 | Intergenic | N/A |
| ctg3568 | 716,102 | 716,174 | 73 | 29.71 | 5 | 5 | < 0.00001 | Intergenic | N/A |
| ctg3738 | 186,066 | 186,201 | 136 | 27.92 | 6 | 3 | 0.00003 | *MMCM3* | Maternal DNA replication licensing factor mcm3 |
| ctg3758**^b^** | 44,460 | 44,677 | 218 | -27.97 | 13 | 9 | < 0.00001 | Intergenic | N/A |
| ctg3759**^a^** | 6,756 | 6,870 | 115 | -38.87 | 7 | 7 | < 0.00001 | Intergenic | N/A |
| ctg3773**^b^** | 120,249 | 120,346 | 98 | -22.77 | 7 | 5 | < 0.00001 | Promoter of *SOCS3* | Suppressor of cytokine signaling 3 |
| ctg3773**^b^** | 120,787 | 121,112 | 326 | -22.58 | 22 | 10 | < 0.00001 | *SOCS3* | Suppressor of cytokine signaling 3 |
| ctg3773**^b^** | 171,406 | 171,542 | 137 | -22.15 | 10 | 5 | 0.00004 | *PGS1* | CDP-diacylglycerol-glycerol-3-phosphate 3-phosphatidyltransferase, mitochondrial |
| ctg4046 | 115,513 | 115,634 | 122 | 26.41 | 6 | 4 | < 0.00001 | *RAB6A* | Ras-related protein Rab-6A |
| ctg42**^a^** | 1,763,592 | 1,763,650 | 59 | 31.94 | 6 | 6 | < 0.00001 | *PKP4* | Plakophilin-4 |
| ctg4628 | 112,210 | 112,354 | 145 | 22.45 | 13 | 4 | 0.00028 | Intergenic | N/A |
| ctg4676 | 16,488 | 16,607 | 120 | -21.78 | 7 | 3 | < 0.00001 | *TCB1* | Transposable element Tcb1 transposase |
| ctg4820 | 179,259 | 179,361 | 103 | 22.70 | 9 | 4 | < 0.00001 | *SKIV2L2* | Superkiller viralicidic activity 2-like 2 |
| ctg5009 | 155,256 | 155,336 | 81 | 31.43 | 6 | 5 | < 0.00001 | Intergenic | N/A |
| ctg505 | 18,210 | 18,349 | 140 | -37.37 | 8 | 8 | < 0.00001 | Intergenic | N/A |
| ctg508 | 90,532 | 90,621 | 90 | 22.70 | 11 | 5 | < 0.00001 | Intergenic | N/A |
| ctg5287 | 289,785 | 289,864 | 80 | -27.96 | 10 | 8 | < 0.00001 | *TRMT61A* | tRNA (adenine(58)-N(1))-methyltransferase catalytic subunit TRMT61A |
| ctg5424**^b^** | 170,718 | 170,848 | 131 | 36.03 | 5 | 4 | < 0.00001 | *HSPB6* | Heat shock protein beta-6 |
| ctg6018 | 72,530 | 72,774 | 245 | 25.33 | 8 | 3 | 0.00005 | Intergenic | N/A |
| ctg6194 | 82,802 | 82,894 | 93 | 31.87 | 8 | 8 | < 0.00001 | *RMA_00027785* | Unknown |
| ctg6434**^b^** | 97,065 | 97,333 | 269 | 21.67 | 10 | 3 | 0.00033 | *SCNN1G* | Amiloride-sensitive sodium channel subunit gamma |
| ctg6628 | 18,146 | 18,198 | 53 | 25.17 | 8 | 3 | 0.00016 | *RMA_00047359* | Unknown |
| ctg6924 | 35,252 | 35,360 | 109 | -20.87 | 6 | 3 | 0.0016 | Intergenic | N/A |
| ctg7054**^a^** | 25,391 | 25,575 | 185 | 29.34 | 11 | 6 | < 0.00001 | Intergenic | N/A |
| ctg7585**^c^** | 41,611 | 41,764 | 154 | 32.81 | 11 | 10 | < 0.00001 | *FAM168A* | Protein FAM168A |
| ctg76 | 43,940 | 44,093 | 154 | 67.00 | 11 | 11 | < 0.00001 | Intergenic | N/A |
| ctg7863**^c^** | 136,569 | 136,654 | 86 | -40.76 | 6 | 5 | < 0.00001 | Intergenic | N/A |
| ctg7875 | 169,559 | 169,767 | 209 | 29.38 | 15 | 5 | < 0.00001 | Intergenic | N/A |
| ctg7948 | 42,453 | 42,704 | 252 | 39.57 | 11 | 8 | < 0.00001 | Intergenic | N/A |
| ctg8148 | 134,671 | 135,196 | 526 | 27.27 | 32 | 14 | < 0.00001 | Intergenic | N/A |
| ctg8148 | 161,076 | 161,507 | 432 | -21.49 | 11 | 3 | 0.00005 | Intergenic | N/A |
| ctg8148 | 161,900 | 162,142 | 243 | -30.57 | 11 | 9 | < 0.00001 | Intergenic | N/A |
| ctg892 | 3,081 | 3,230 | 150 | -29.49 | 5 | 3 | 0.00003 | Intergenic | N/A |
| ctg913 | 1,182,605 | 1,183,295 | 691 | 21.84 | 16 | 3 | 0.0013 | *NLK* | Serine/threonine-protein kinase NLK |
| ctg994**^a^** | 348,235 | 348,448 | 214 | 43.47 | 8 | 6 | < 0.00001 | Intergenic | N/A |
| ctg998**^c^** | 35,687 | 35,889 | 203 | -21.16 | 7 | 3 | 0.00001 | *HYDIN* | Hydrocephalus-inducing protein homolog |

**^a^**Also found between range-core tadpoles exposed to alarm cues and controls.

**^b^**Also found between range-edge tadpoles exposed to alarm cues and controls.

**^c^**Also found between range-core tadpoles exposed to cannibal cues and controls.

**Table S4.** DMRs between range-core tadpoles exposed to alarm cues and controls.

| Contig | Start DMR | End DMR | Length DMR (bp) | Methylation difference (%) | No. CpGs | No. DMCs | DMR  *q*-value | Gene | Protein |
| --- | --- | --- | --- | --- | --- | --- | --- | --- | --- |
| ctg10122 | 21,047 | 21,112 | 66 | -25.03 | 5 | 3 | 0.0012 | *MYL3* | Myosin light chain 3 |
| ctg10322 | 63,251 | 63,313 | 63 | -21.67 | 6 | 3 | < 0.00001 | *PLA2G4A* | Cytosolic phospholipase A2 |
| ctg11043 | 174,618 | 174,666 | 49 | 32.98 | 7 | 3 | < 0.00001 | Intergenic | N/A |
| ctg12425 | 176,373 | 176,426 | 54 | 32.97 | 5 | 4 | < 0.00001 | *ARHGAP9* | Rho GTPase-activating protein 9 |
| ctg1295 | 229,245 | 229,311 | 67 | 23.13 | 5 | 4 | < 0.00001 | Promoter of *HGSNAT* | Heparan-alpha-glucosaminide N-acetyltransferase |
| ctg13303 | 30,499 | 30,669 | 171 | 23.70 | 11 | 7 | < 0.00001 | *RMA_00047991* | Unknown |
| ctg13763 | 5,234 | 5,360 | 127 | 25.48 | 5 | 4 | < 0.00001 | Intergenic | N/A |
| ctg14203 | 43,476 | 43,558 | 83 | 39.58 | 6 | 5 | < 0.00001 | *NXPE4* | NXPE family member 4 |
| ctg1466 | 150,552 | 150,774 | 223 | 20.58 | 14 | 4 | 0.00013 | *EIF3A* | Eukaryotic translation initiation factor 3 subunit A |
| ctg1550 | 233,177 | 233,268 | 92 | 24.80 | 8 | 4 | 0.00002 | Intergenic | N/A |
| ctg17636 | 94,671 | 94,908 | 238 | 22.36 | 18 | 15 | < 0.00001 | *ZFAND3* | AN1-type zinc finger protein 3 homolog |
| ctg1915 | 401,724 | 401,823 | 100 | -24.24 | 7 | 4 | < 0.00001 | Promoter of *PITPNC1* | Cytoplasmic phosphatidylinositol transfer protein 1 |
| ctg19786**^c^** | 63,198 | 63,623 | 426 | -21.14 | 15 | 6 | < 0.00001 | Intergenic | N/A |
| ctg20587 | 198,837 | 198,978 | 142 | 25.20 | 6 | 4 | < 0.00001 | Intergenic | N/A |
| ctg23772 | 30,847 | 30,936 | 90 | -26.49 | 7 | 5 | < 0.00001 | Promoter of *ST6GALNAC4* | Alpha-N-acetyl-neuraminyl-2,3-beta-galactosyl-1,3-N-acetyl-galactosaminide alpha-2,6-sialyltransferase |
| ctg2404 | 71,060 | 71,357 | 298 | -24.22 | 5 | 3 | 0.00006 | *CEP135* | Centrosomal protein of 135 kDa |
| ctg24887 | 9,171 | 9,211 | 41 | 24.56 | 5 | 3 | 0.00002 | Intergenic | N/A |
| ctg29360 | 6,709 | 6,782 | 74 | 22.09 | 6 | 3 | 0.00013 | Intergenic | N/A |
| ctg3020 | 75,379 | 75,555 | 177 | -30.32 | 8 | 4 | < 0.00001 | *RNASEH1* | Ribonuclease H1 |
| ctg3373 | 115,262 | 115,387 | 126 | 23.44 | 5 | 3 | < 0.00001 | Intergenic | N/A |
| ctg3633 | 303,843 | 304,055 | 213 | 26.17 | 9 | 4 | 0.00053 | Intergenic | N/A |
| ctg3759**^c^** | 6,756 | 6,870 | 115 | -20.06 | 7 | 4 | < 0.00001 | Intergenic | N/A |
| ctg3763 | 35,833 | 36,011 | 179 | 24.78 | 11 | 7 | < 0.00001 | Intergenic | N/A |
| ctg3915 | 764,591 | 764,690 | 100 | 21.54 | 8 | 5 | < 0.00001 | *GPR155* | Integral membrane protein GPR155 |
| ctg42**^c^** | 1,763,592 | 1,763,650 | 59 | -24.50 | 6 | 5 | < 0.00001 | *PKP4* | Plakophilin-4 |
| ctg4301 | 45,500 | 45,663 | 164 | 32.95 | 7 | 5 | < 0.00001 | *ACYP2* | Acylphosphatase-2 |
| ctg4767 | 166,936 | 167,026 | 91 | 57.13 | 7 | 7 | < 0.00001 | *COX10* | Protoheme IX farnesyltransferase, mitochondrial |
| ctg547 | 256,202 | 256,535 | 334 | 29.59 | 6 | 4 | < 0.00001 | Intergenic | N/A |
| ctg5501 | 230,535 | 230,708 | 174 | 26.15 | 19 | 6 | 0.00003 | *DACH2* | Dachshund homolog 2 |
| ctg552 | 340,264 | 340,358 | 95 | 27.93 | 7 | 6 | < 0.00001 | *GRAP* | GRB2-related adapter protein |
| ctg561 | 74,350 | 74,574 | 225 | 43.10 | 15 | 11 | < 0.00001 | Intergenic | N/A |
| ctg56 | 217,365 | 217,478 | 114 | 20.74 | 6 | 3 | 0.00054 | *ARIH2* | E3 ubiquitin-protein ligase ARIH2 |
| ctg6147 | 93,558 | 94,000 | 443 | -25.75 | 13 | 12 | < 0.00001 | *CFAP77* | Cilia- and flagella-associated protein 77 |
| ctg6147 | 260,373 | 260,587 | 215 | -29.43 | 14 | 10 | < 0.00001 | Intergenic | N/A |
| ctg6543 | 122,891 | 123,041 | 151 | 20.25 | 7 | 4 | < 0.00001 | Intergenic | N/A |
| ctg7054**^c^** | 25,391 | 25,575 | 185 | 33.29 | 11 | 6 | < 0.00001 | Intergenic | N/A |
| ctg7892 | 85,601 | 85,663 | 63 | 20.04 | 5 | 3 | < 0.00001 | Promoter of *RMA_00030658* | Unknown |
| ctg8084**^a^** | 148,657 | 148,744 | 88 | 26.62 | 5 | 3 | < 0.00001 | *GUCA1A* | Guanylyl cyclase-activating protein 1 |
| ctg8225 | 102,294 | 102,357 | 64 | 30.75 | 8 | 3 | < 0.00001 | *ENG* | Endoglin |
| ctg834 | 307,186 | 307,251 | 66 | 23.87 | 5 | 4 | < 0.00001 | Intergenic | N/A |
| ctg9382 | 50,247 | 50,337 | 91 | 27.01 | 8 | 6 | < 0.00001 | Intergenic | N/A |
| ctg994**^c^** | 348,001 | 348,448 | 448 | -21.81 | 16 | 7 | < 0.00001 | Intergenic | N/A |
| ctg9969**^b^** | 28,668 | 28,807 | 140 | 27.05 | 5 | 3 | < 0.00001 | Promoter of *RMA_00030658* | Unknown |

**^a^**Also found between range-edge tadpoles exposed to alarm cues and controls.

**^b^**Also found between range-core tadpoles exposed to cannibal cues and controls.

**^c^**Also found between range-edge tadpoles exposed to cannibal cues and controls.

**Table S5.** DMRs between range-edge tadpoles exposed to alarm cues and controls.

| Contig | Start DMR | End DMR | Length DMR (bp) | Methylation difference (%) | No. CpGs | No. DMCs | DMR  *q*-value | Gene | Protein |
| --- | --- | --- | --- | --- | --- | --- | --- | --- | --- |
| ctg1044 | 3,039 | 3,347 | 309 | 23.47 | 11 | 10 | < 0.00001 | *CD63* | CD63 antigen |
| ctg111 | 23,206 | 23,369 | 164 | -28.05 | 5 | 4 | < 0.00001 | Intergenic | N/A |
| ctg1181 | 1,649 | 1,851 | 203 | 21.44 | 12 | 6 | < 0.00001 | *FCF1* | rRNA-processing protein FCF1 homolog |
| ctg1181**^c^** | 2,961 | 3,382 | 422 | 22.90 | 16 | 10 | < 0.00001 | Intergenic | N/A |
| ctg1325 | 156,402 | 156,483 | 82 | 27.99 | 6 | 4 | < 0.00001 | *PRKCZ* | Protein kinase C zeta type |
| ctg1428 | 238,871 | 238,986 | 116 | 21.59 | 6 | 3 | < 0.00001 | *TIE1* | Tyrosine-protein kinase receptor Tie-1 |
| ctg1543 | 245,121 | 245,213 | 93 | 38.13 | 5 | 3 | < 0.00001 | *L1RE1* | LINE-1 retrotransposable element ORF1 protein |
| ctg1563 | 762,370 | 762,895 | 526 | -20.97 | 11 | 6 | < 0.00001 | Intergenic | N/A |
| ctg1699 | 27,371 | 27,721 | 351 | -24.68 | 9 | 4 | < 0.00001 | Intergenic | N/A |
| ctg17660 | 14,354 | 14,430 | 77 | 21.85 | 7 | 6 | < 0.00001 | Intergenic | N/A |
| ctg17928 | 59,148 | 59,241 | 94 | 29.58 | 9 | 8 | < 0.00001 | Intergenic | N/A |
| ctg18198 | 77,767 | 78,023 | 257 | 26.07 | 17 | 6 | < 0.00001 | *PHYH* | Phytanoyl-CoA dioxygenase, peroxisomal |
| ctg1899 | 133,489 | 133,541 | 53 | 20.57 | 5 | 4 | < 0.00001 | *MST1R* | Macrophage-stimulating protein receptor |
| ctg19321**^c^** | 2,398 | 2,512 | 115 | 21.44 | 7 | 5 | < 0.00001 | *RMA_00054127* | Unknown |
| ctg21885 | 14,026 | 14,189 | 164 | 46.15 | 9 | 9 | < 0.00001 | *ZCCHC3* | Zinc finger CCHC domain-containing protein 3 |
| ctg21885 | 15,781 | 15,992 | 212 | 45.18 | 5 | 5 | < 0.00001 | *ZCCHC3* | Zinc finger CCHC domain-containing protein 3 |
| ctg21885 | 16,381 | 16,960 | 580 | 42.73 | 25 | 25 | < 0.00001 | *ZCCHC3* | Zinc finger CCHC domain-containing protein 3 |
| ctg21885 | 17,460 | 17,545 | 86 | 32.86 | 7 | 5 | < 0.00001 | *ZCCHC3* | Zinc finger CCHC domain-containing protein 3 |
| ctg21994 | 18,763 | 18,920 | 158 | -36.07 | 11 | 9 | < 0.00001 | Intergenic | N/A |
| ctg22378 | 15,218 | 15,321 | 104 | 26.00 | 8 | 5 | < 0.00001 | Intergenic | N/A |
| ctg24612 | 52,646 | 52,945 | 300 | 21.05 | 20 | 11 | < 0.00001 | Intergenic | N/A |
| ctg2502 | 121,898 | 121,990 | 93 | 36.61 | 5 | 5 | < 0.00001 | Intergenic | N/A |
| ctg2506 | 228,750 | 228,816 | 67 | 30.41 | 6 | 3 | < 0.00001 | Promoter of *ECE1* | Endothelin-converting enzyme 1 |
| ctg25970**^c^** | 30,960 | 31,112 | 153 | 37.25 | 13 | 13 | < 0.00001 | Intergenic | N/A |
| ctg26366 | 16,221 | 16,386 | 166 | 30.65 | 6 | 4 | < 0.00001 | Intergenic | N/A |
| ctg2771 | 629,305 | 629,399 | 95 | -28.13 | 8 | 4 | 0.00043 | *HPCAL1* | Hippocalcin-like protein 1 |
| ctg2853 | 338,699 | 338,786 | 88 | -26.29 | 5 | 4 | < 0.00001 | *RMA_00013077* | Unknown |
| ctg2982 | 394,793 | 394,914 | 122 | -23.83 | 10 | 8 | < 0.00001 | *RIMS2* | Regulating synaptic membrane exocytosis protein 2 |
| ctg2989 | 8,014 | 8,094 | 81 | -43.56 | 5 | 5 | < 0.00001 | *KCNK10* | Potassium channel subfamily K member 10 |
| ctg30982 | 9,910 | 9,982 | 73 | 23.30 | 8 | 6 | < 0.00001 | Intergenic | N/A |
| ctg3616 | 109,281 | 109,463 | 183 | 21.72 | 8 | 3 | 0.00010 | Intergenic | N/A |
| ctg3705 | 442,058 | 442,262 | 205 | 35.97 | 7 | 5 | < 0.00001 | Intergenic | N/A |
| ctg3707 | 72,464 | 72,610 | 147 | 21.45 | 9 | 5 | < 0.00001 | Intergenic | N/A |
| ctg3758**^c^** | 44,460 | 44,677 | 218 | 23.35 | 13 | 7 | < 0.00001 | Intergenic | N/A |
| ctg3773**^c^** | 120,179 | 120,346 | 168 | -21.59 | 11 | 6 | < 0.00001 | Promoter of *SOCS3* | Suppressor of cytokine signaling 3 |
| ctg3773**^c^** | 120,787 | 121,180 | 394 | -34.95 | 25 | 21 | < 0.00001 | *SOCS3* | Suppressor of cytokine signaling 3 |
| ctg3773**^c^** | 171,024 | 171,542 | 519 | -21.52 | 15 | 6 | < 0.00001 | *PGS1* | CDP-diacylglycerol--glycerol-3-phosphate 3-phosphatidyltransferase, mitochondrial |
| ctg399**^b^** | 944,763 | 945,066 | 304 | 29.24 | 19 | 12 | < 0.00001 | *RMA_00000723* | Unknown |
| ctg4038 | 34,701 | 34,793 | 93 | 20.37 | 7 | 4 | < 0.00001 | Intergenic | N/A |
| ctg4124 | 248,589 | 248,766 | 178 | 35.49 | 8 | 5 | < 0.00001 | *PARD6B* | Partitioning defective 6 homolog beta |
| ctg5424**^c^** | 170,718 | 170,848 | 131 | -27.62 | 5 | 4 | < 0.00001 | *HSPB6* | Heat shock protein beta-6 |
| ctg5497 | 14,077 | 14,209 | 133 | 24.35 | 5 | 3 | 0.00005 | *C11ORF96* | Uncharacterized protein C11orf96 |
| ctg5507 | 2,222 | 2,483 | 262 | -21.88 | 11 | 8 | < 0.00001 | Intergenic | N/A |
| ctg6147 | 375,208 | 375,342 | 135 | -22.06 | 12 | 5 | < 0.00001 | Intergenic | N/A |
| ctg6344 | 100,935 | 101,066 | 132 | -27.92 | 14 | 5 | 0.00004 | *EPHB5* | Ephrin type-B receptor 5 |
| ctg6434**^c^** | 97,065 | 97,333 | 269 | 20.82 | 10 | 8 | < 0.00001 | *SCNN1G* | Amiloride-sensitive sodium channel subunit gamma |
| ctg6434 | 98,765 | 98,965 | 201 | -25.83 | 8 | 3 | < 0.00001 | *SCNN1G* | Amiloride-sensitive sodium channel subunit gamma |
| ctg6643 | 54,137 | 54,327 | 191 | 28.76 | 14 | 10 | < 0.00001 | Intergenic | N/A |
| ctg6922 | 112,733 | 112,867 | 135 | 52.82 | 5 | 4 | < 0.00001 | Intergenic | N/A |
| ctg8084**^a^** | 148,657 | 148,744 | 88 | 28.43 | 5 | 3 | < 0.00001 | *GUCA1A* | Guanylyl cyclase-activating protein 1 |
| ctg8281 | 36,970 | 37,033 | 64 | -25.67 | 7 | 7 | < 0.00001 | Intergenic | N/A |
| ctg9467 | 108,460 | 108,672 | 213 | -24.32 | 5 | 3 | 0.0081 | *MINK1* | Misshapen-like kinase 1 |
| ctg966 | 144,472 | 144,685 | 214 | 22.53 | 12 | 11 | < 0.00001 | Intergenic | N/A |

**^a^**Also found between range-core tadpoles exposed to alarm cues and controls.

**^b^**Also found between range-core tadpoles exposed to cannibal cues and controls.

**^c^**Also found between range-edge tadpoles exposed to cannibal cues and controls.
